# Dynamic Subcortical Modulators of Human Default Mode Network Function

**DOI:** 10.1101/2021.10.27.466172

**Authors:** Ben J. Harrison, Christopher G. Davey, Hannah S. Savage, Alec J. Jamieson, Christine A. Leonards, Bradford A. Moffat, Rebecca K. Glarin, Trevor Steward

## Abstract

The brain’s ‘default mode network’ (DMN) enables flexible switching between internally and externally focused cognition. Precisely how this modulation occurs is not well understood, although may involve key subcortical mechanisms, including hypothesized influences from the basal forebrain (BF) and mediodorsal thalamus (MD). Here, we used ultra-high field (7T) functional magnetic resonance imaging to examine the involvement of the BF and MD across states of task-induced DMN activity modulation. Specifically, we mapped DMN activity suppression (‘deactivation’) when participants transitioned between rest and externally focused task performance, as well as DMN activity engagement (‘activation’) when task performance was internally (i.e., self) focused. Consistent with recent rodent studies, the BF showed overall activity suppression with DMN cortical regions when comparing the rest to external task conditions. Further analyses, including dynamic causal modelling, confirmed that the BF drove changes in DMN cortical activity during these rest-to-task transitions. The MD, by comparison, was specifically engaged during internally focused cognition and demonstrated a broad excitatory influence on DMN cortical activation. These results provide the first direct evidence in humans of distinct basal forebrain and thalamic circuit influences on the control of DMN function and suggest novel mechanistic avenues for ongoing translational research.

## Introduction

Since its first characterization over two decades ago (Andreasen et al. 1995; Shulman et al. 1997; Raichle et al. 2001), the brain’s ‘default mode network’ (DMN) has achieved a special status in the neuroscientific study of cognition and behavior (Davey and Harrison 2018; Buckner and DiNicola 2019; Yeshurun et al. 2021). Current conceptualizations of the DMN, for instance, place it atop of a rich hierarchy of brain systems supporting advanced cognitive processes – a view that emphasizes its unique anatomical organization and connectivity (Margulies et al. 2016; Smallwood et al. 2021). DMN function, in particular, has been predominantly linked to internally-driven cognitive modes, such as self-directed thought, which rely on internal mental representations rather than immediate processing of the external environment (Gusnard et al. 2001; Buckner and Carroll 2007; Andrews-Hanna et al. 2014; Davey and Harrison 2018). Precisely how the DMN coordinates these processes is not yet well understood, though must involve mechanisms that enable the control and integration of activity across its widespread cortical territories, spanning the frontal and posterior midline, and inferior parietal cortices. Based on emerging evidence from human imaging studies, as well as other novel insights (Nair et al. 2018), there is reason to believe that such mechanisms might exist in deeper subcortical structures of the brain, despite such structures being traditionally overlooked in the study of the DMN.

In important recent work, Alves and colleagues (2019) provided an optimized mapping of DMN structural and functional connectivity that identified the basal forebrain and anterior- and mediodorsal thalamus as its major subcortical nodes, with evidence to suggest specific roles in network integration and resilience. Their findings with respect to the basal forebrain (BF) complement other recent imaging studies that have mapped its functional connectivity with DMN cortical regions (Markello et al. 2018; Yuan et al. 2019), as well as evidence from rodents indicating a role for the BF in modulating DMN dynamics (Nair et al. 2018; Lozano-Montes et al. 2020). In this latter work, BF neurons were shown to exhibit robust activity suppression when animals were transitioned between resting-state and novel task environments – a result that mirrors the well-known pattern of DMN suppression (‘deactivation’) in humans when performing externally-focused imaging tasks (Shulman et al. 1997; Raichle et al. 2001; Harrison et al. 2008). Critically, in the rodent studies, BF signaling was further shown to drive changes in medial frontal cortex activity, especially during rest, suggesting a mechanism for regulating the DMN between internal and external cognitive states (Nair et al. 2018). Thus, while deactivation has been long appreciated as a unique characteristic of the DMN that reflects its role in facilitating cognitive state-transitions, it is only recently that specific mechanisms underlying this phenomenon have been examined.

By comparison, renewed interest in the contribution of the thalamus to higher cognition focuses on its hypothesized role in coordinating and sustaining information flow and plasticity across large-scale cortical networks (Parnaudeau et al. 2018; Pergola et al. 2018; Halassa and Sherman 2019; Wolff and Vann 2019). Of the higher-order thalamic nuclei, the mediodorsal thalamus (MD) shows preferential connectivity with midline frontal regions of the DMN, especially the medial prefrontal cortex (MPFC), whereas the anterior thalamus is more densely connected to its posterior midline (Goldman-Rakic and Porrino 1985; Vogt et al. 1987; Barbas 2000; Aggleton et al. 2014; Phillips et al. 2019). Both divisions were implicated as DMN hubs in the Alves et al. (2019) study, whereas in a related study by Greene et al. (2020), the MD was mapped more distinctly as a ‘cognitive integration zone’, linking the DMN to other large-scale cortical networks. Recent work by our group has complemented these findings by mapping a broad excitatory influence of the MD on multiple prefrontal regions, including the MPFC, during a cognitive self-regulation task (Steward et al. 2021). Its results argue for a distinct coordinating role of the MD during higher cognition, whereby its task-evoked activity helps to sustain the engagement of cortical function.

In the current study, we set out to examine the roles of the BF and MD on the function of DMN cortical regions, hypothesizing distinct modulatory influences during states of task-induced deactivation and activation, respectively. Specifically, we hypothesized that the BF would be involved in modulating DMN cortical activity between rest and external cognitive states, consistent with its role in driving DMN state transitions, whereas the MD would broadly upregulate DMN cortical activity during self-oriented higher cognition. The task chosen to address these aims was used in previous work by our group that established a dynamic network model of DMN function, targeting its primary cortical regions, the MPFC, posterior cingulate cortex (PCC) and inferior parietal lobule (IPL; Davey et al. 2016; Davey et al. 2017). These past results also support the task’s capacity to induce significant deactivation of the BF when external cognition is compared to rest, and activation of the MD during self-related versus external cognition (Davey et al. 2016). Here, we used ultra-high field functional magnetic resonance imaging (UHF fMRI) to target their involvement more precisely and to test neural models of their directional influence on DMN cortical regions, with results confirming their unique contributions to network functioning.

## Materials and Methods

### Participants

Forty participants were recruited to the study. All participants met the following eligibility criteria: (i) they were aged between 18 and 25 years; (ii) had no current or past diagnosis of mental illness; (iii) were competent English speakers, (iv) were not taking any psychoactive medication; and (v) had no contraindications to MRI, including pregnancy. All participants had normal or corrected-to-normal vision and provided written informed consent, following a complete description of the study protocol, which was approved by The University of Melbourne Human Research Ethics Committee. Of the initial sample, 2 participants did not complete the task; 1 did not have physiological data available; and a further 3 participants were excluded due to excessive head motion (see ‘Image Pre-processing’). The final sample consisted of 34 participants (21 female) with a mean age of 22.0 years (± 2.0 years).

### Experimental Design

Participants completed the ‘self-appraisal’ task, as detailed in Davey et al., (2016), which comprised three task elements: self-related cognition, external cognition, and rest. In the self and external conditions, participants were presented with personality trait adjectives and responded to them as instructed below. Words were drawn from a frequently used list of trait adjectives (Anderson 1968): we selected 96 words distributed around the median rating for ‘likeableness’ reported in the original dataset. The words were selected so as not to be extremely favorable or unfavorable, with the intention of engaging participants in self-referential appraisal (adjectives included words such as ‘skeptical’, ‘perfectionistic’, and ‘lucky’). For the self condition, participants viewed 8 blocks of 6 words, presented for 5 sec each, and responded to the question, ‘*Does this word describe you?*’, by pressing the left or right button on the button-box (Lumina, Cedrus Corporation). In the external condition, participants viewed 8 blocks of 6 words, also presented for 5 s each, and responded to the question, ‘*Does this word have 4 or more vowels?* ‘. The 2 lists of 48 words that formed these conditions were matched on valence and number of vowels and were counterbalanced across participants. Each 32 s block (2 s instruction followed by 6 words presented for 5 s each) was interspersed with a 10 s rest-fixation block (17 in total) in which participants were asked to fixate on a centrally presented cross-hair. The task was programmed in Paradigm software (http://www.paradigmexperiments.com) and was presented on a 32” LCD BOLD screen (Cambridge Research Systems) visible via a reverse mirror mounted to the head coil.

### UHF Image Acquisition

Imaging was performed on a 7T research scanner (Siemens Healthcare, Erlangen, Germany) equipped with a 32-channel head-coil (Nova Medical Inc., Wilmington MA, USA). The functional sequence consisted of a multi-band (6 times) and grappa (2 times) accelerated GE-EPI sequence in the steady state (TR, 800ms; TE, 22.2 ms; pulse/flip angle, 45°; field-of-view, 20.8 cm; slice thickness (no gap), 1.6 mm; 130 × 130-pixel matrix; 84 interleaved axial slices aligned to AC-PC line; Setsompop et al. 2012). The total sequence time was 11 minutes and 22 seconds, corresponding to 852 whole-brain EPI volumes. A T1-weighted high-resolution anatomical image (MP2RAGE; Marques et al. 2010) was acquired for each participant to assist with functional time series co-registration (TR = 5000 ms; TE, 3.0 ms; inversion times, 700/2700ms; pulse/flip angles, 4/5°; field-of-view, 24 cm; slice thickness (no gap), 0.73 mm; 330 × 330–pixel matrix; 84 sagittal slices aligned parallel to the midline). The total sequence time was 7 minutes and 12 seconds. To assist with head immobility, foam-padding inserts were placed on either side of the participants’ head. Cardiac and respiratory recordings were sampled at 50 Hertz (Hz) using a Siemens (Bluetooth) pulse-oximeter and respiratory belt. Information derived from these recordings were used for physiological noise correction (see further).

### Image Pre-processing

Imaging data were pre-processed using Statistical Parametric Mapping (SPM) 12 (v7771, Welcome Trust Centre for Neuroimaging, London, UK) within a MATLAB 2019b environment (The MathWorks Inc., Natick, MA). Motion artifacts were corrected by realigning each participant’s time-series to the mean image, and all images were resampled using 4th Degree B-Spline interpolation. Individualized motion regressors were created using the Motion Fingerprint approach to account for movement (Wilke 2012). As noted above, 3 participants were excluded due to a mean total scan-to-scan displacement over 1.6 mm (i.e., the size of one voxel). Each participant’s anatomical images were co-registered to their respective mean functional image, segmented and normalized to the International Consortium of Brain Mapping template using the unified segmentation plus DARTEL approach. Smoothing was applied with a 3.2 mm^3^ full-width-at-half-maximum (FWHM) Gaussian kernel to preserve spatial specificity.

Physiological noise was modelled at the first level using the PhysIO Toolbox (Kasper et al. 2017). This toolbox applies noise correction to fMRI sequences using physiological recordings and has been found to improve blood-oxygen level dependent (BOLD) signal sensitivity and temporal signal-to-noise ratio (tSNR) at 7T (Reynaud et al. 2017; see also Steward et al. 2021). The Retrospective Image-based Correction function (RETROICOR; Glover et al. 2000) was applied to model the periodic effects of heartbeat and breathing on BOLD signals, using acquired cardiac/respiratory phase information. The Respiratory Response Function (RRF; Birn et al. 2008), convolved with respiration volume per time (RVT) was used to model low frequency signal fluctuations, which arose from changes in breathing depth and rate. Heart rate variability (HRV) was convolved with a predefined Cardiac Response Function (CRF; Chang et al. 2009) to account for BOLD variances due to heartrate-dependent changes in blood oxygenation.

### General Linear Modelling

Each participant’s pre-processed time-series and nuisance regressors (i.e., physiological noise and Motion Fingerprint regressors) were included in a first level GLM analysis. Primary regressors for the self and external conditions, as well as task instruction periods, were created by specifying the onset and duration of each block, followed by convolution with a canonical hemodynamic response function. The rest condition periods formed an implicit baseline. A 128-Hz high-pass filter was applied to account for low-frequency noise and temporal autocorrelation was estimated using SPM’s FAST method, which has been shown to outperform AR(1) at short TRs and yield superior reliability (Olszowy et al. 2019). Primary contrast images (external vs. baseline/rest; self vs baseline/rest) for each participant were entered in a second-level random-effects GLM analysis (2 level, within-subjects ANOVA). Three main group effects were estimated from this model: rest > external (to map regions of DMN deactivation); self > external (to map regions of DMN activation); and their conjunction (to map common ‘core’ DMN cortical regions; see Davey et al. 2016). These analyses were used as both initial tests of our hypotheses regarding the involvement of the BF and MD during states of DMN deactivation and activation, respectively, and to support the mapping of specific regions to be included in dynamic causal modelling (DCM), as will be described below. For all GLM analyses, whole-brain, false discovery rate (FDR) corrected statistical thresholds were applied (*P*_FDR_ <0.05; 10 voxel cluster-extent threshold).

### Dynamic Causal Modeling

The aim of dynamic causal modeling (DCM) is to infer the causal architecture of a network of brain regions, which in the case of fMRI time-series data, involves using generative models to estimate functional interactions from underlying (hidden) neurophysiological activity (Friston et al. 2003; Stephan et al. 2010; Friston et al. 2019). Using a Bayesian approach, a model is selected from a set of predefined models that is the most likely to generate the observed imaging data while also penalizing for model complexity. DCMs delineate how dynamics in one brain region influence dynamics in others, incorporating both the core set of interregional (‘baseline’) connections and the modulation of those influences by experimental (i.e., task) manipulation. As outlined in Stephen et al. (2010), DCMs are both sensitive to temporal precedence effects between neuronal populations (i.e., how the present activity state of one region causes change in other), as well as the spatiotemporal structure of a network’s dynamics: that is, specifically ‘where and when’ these interactions occur under the influence of experimental (i.e., task) manipulations. The relative evidence of these causal influences can then be compared through model comparison to determine which model architecture optimally explains the data. Compared to temporal correlation measures of functional connectivity, DCM provides a more detailed and physiologically valid mapping of effective connectivity – the directed causal influences of brain regions on one another. In DCM, modulation is measured in Hz, which signifies the rates of change in activity (connection strength) caused by the influence of one region on another. Positive effective connectivity reflects putative excitatory upregulation of activity, whereas negative connectivity represents inhibitory downregulation. Notably, it has been shown that 7T fMRI furnishes more efficient estimates of effective connectivity than those provided at lower field strengths (Tak et al. 2018).

#### Region Selection and Time-Series Extraction

As is standard with DCM, initial group-level GLM analyses were performed to map our specific regions of interest (BF, MD, MPFC, PCC, IPL; see ‘Results’ for a full description). The regional time-series (VOIs) for each of them were extracted at an individual subject-level following recently published guidelines (Zeidman et al. 2019a). This method summarizes regional time-series by calculating the first eigenvariate across all activated voxels within a 4mm radial sphere around the subject specific maxima (p < 0.05, uncorrected), which can be no more than 8 mm from the group maxima. To ensure that voxels included in BF and MD VOIs did not capture adjacent regions, their time-series extraction was run after applying inclusive BF and thalamic masks, respectively, as provided by Alves et al. (2019; https://neurovault.org/collections/CTTXXAYJ/). Because the Alves et al. ‘limbic thalamus’ mask extends to the anterior thalamic nucleus, we further applied the Automated Anatomical Labelling 3 (AAL3) atlas (1×1×1 mm^3^ resolution) to limit data extraction to the MD (Rolls et al. 2020). Participant time-series were pre-whitened to reduce serial correlations, high-pass filtered, and nuisance effects not covered by the ‘effects of interest’ F-contrast were regressed out (i.e., ‘adjusted’ to the F-contrast).

#### Model Specification and Selection

Our model space was specified using DCM 12.5. To address our hypotheses of distinct modulatory influences of the BF and MD, we specified separate full models assuming bidirectional endogenous (‘baseline’) connections between all regions, with the condition ‘task’ – the onset of all blocks comprising the self and rest conditions – set as their driving input. We specified the models separately to prevent unnecessary complexity and to provide the most parsimonious account of the observed model evidence (Stephan et al. 2010). For the BF model, the rest condition was specified as the modulatory input on each connection to and from the BF (i.e., BF *to* MPFC, PCC, IPL, MD and MPFC, PCC, IPL, MD *to* the BF), whereas the self condition was set as the modulatory input for the MD model (MD *to* MPFC, PCC, IPL, MD and MPFC, PCC, IPL, BF *to* MD). In both models, we examined the modulation of BF–MD connections given their well characterized anatomical relationship, particularly regarding BF modulation of MD function (Young et al. 1984; Groenewegen 1988). Figure 1 illustrates the model testing structure for the ‘BF-rest’ and ‘MD-self’ DCMs.

**Figure 1.**
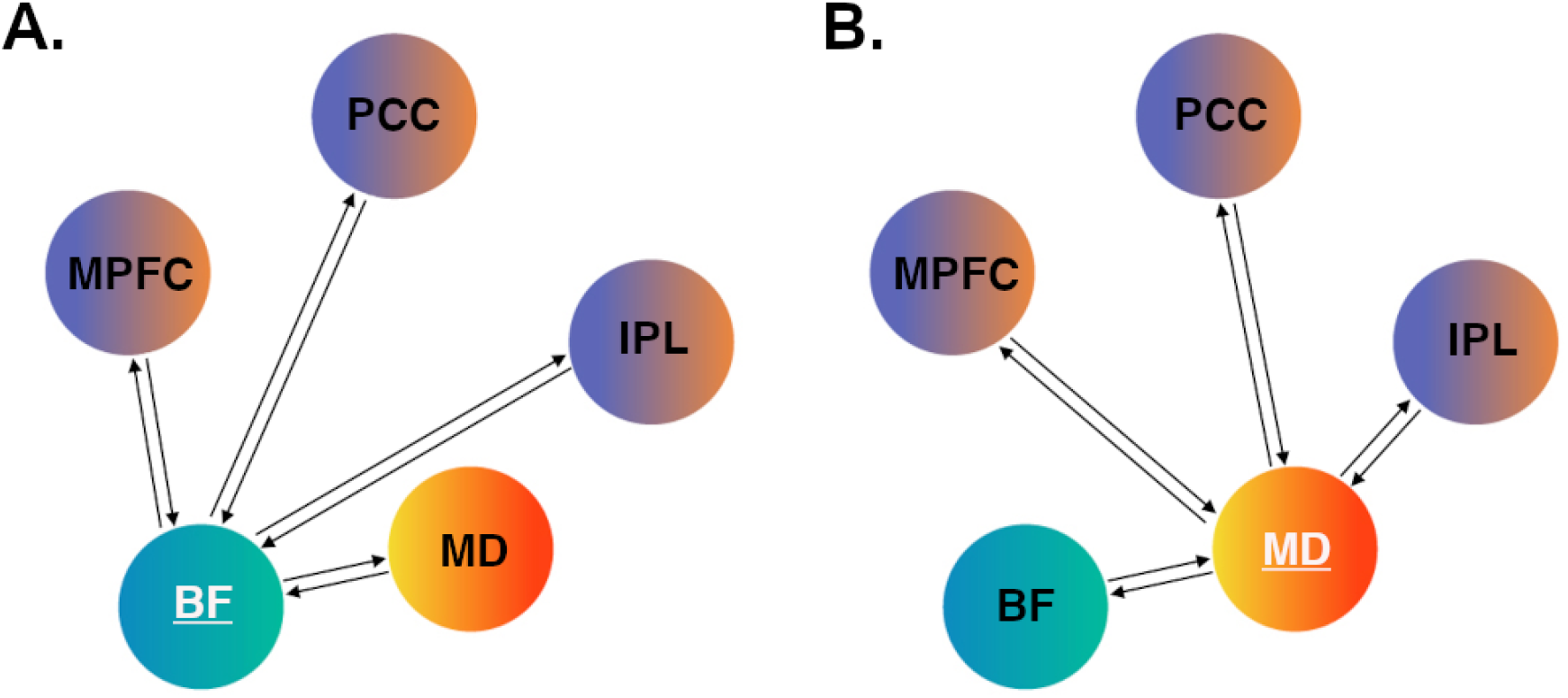
DCM hypothesis testing. A) Basal forebrain (BF) model space; B) Mediodorsal thalamus (MD) model space. For the subcortical region-of-interest in each model (BF or MD), we specified bidirectional intrinsic (baseline) and modulatory connections with the other four regions. For the BF model, modulatory connections were tested under the specific influence of the rest condition. For the MD model, modulatory connections were tested under the specific influence of the self condition. MPFC = medial prefrontal cortex; PCC = posterior cingulate cortex; IPL = inferior parietal lobule.

Full models of effective connectivity were fitted to each participant’s time-series data to calculate the posterior parameter estimates and their probabilities for each connection. At the group level, the connectivity parameter estimates from all participants’ DCMs were assessed using Parametric Empirical Bayes (PEB). The PEB framework affords robust group-level analyses of effective connectivity by means of a hierarchical model, comprising DCMs at the single-subject level and a GLM of connectivity parameters between subjects (Zeidman et al. 2019b). After estimating the PEB model, parameters that did not contribute to the model evidence were pruned using Bayesian model reduction. Bayesian model averaging was then performed over these reduced models to determine parameter estimates: a process which prioritizes simpler and more generalizable models at the group level. A conservative threshold of posterior probability (Pp) > .99 was applied to identify only those parameters which demonstrated very strong evidence (that is, those parameters with a 99% probability of parameters being present vs. absent).

## Results

### General Linear Modelling

As expected, DMN deactivation (rest > external) was associated with broad involvement of characteristic cortical regions of the DMN: the anterior medial wall cortex, spanning ventral to dorsal MPFC, including rostral anterior cingulate cortex; the posteromedial wall cortex, spanning the PCC, retrosplenial, mid cingulate cortex and extending to the precuneus and medial parieto-occipital zone (Figure 2A; Table S1). Additional cortical deactivations included the ventral and dorsal posterior insular cortex; dorsal prefrontal cortex (frontal eye fields); anterior lingual gyrus; anterior and posterior hippocampus (CA1-3; para/subiculum; parahippocampal areas). Subcortically, significant deactivation was observed in the BF; the MD and superior colliculus. Deactivation of the BF was primarily localized to the medial septum and diagonal band areas but extended to surrounding territories including the ventral pallidum (Figure S1A).

**Figure 2.**
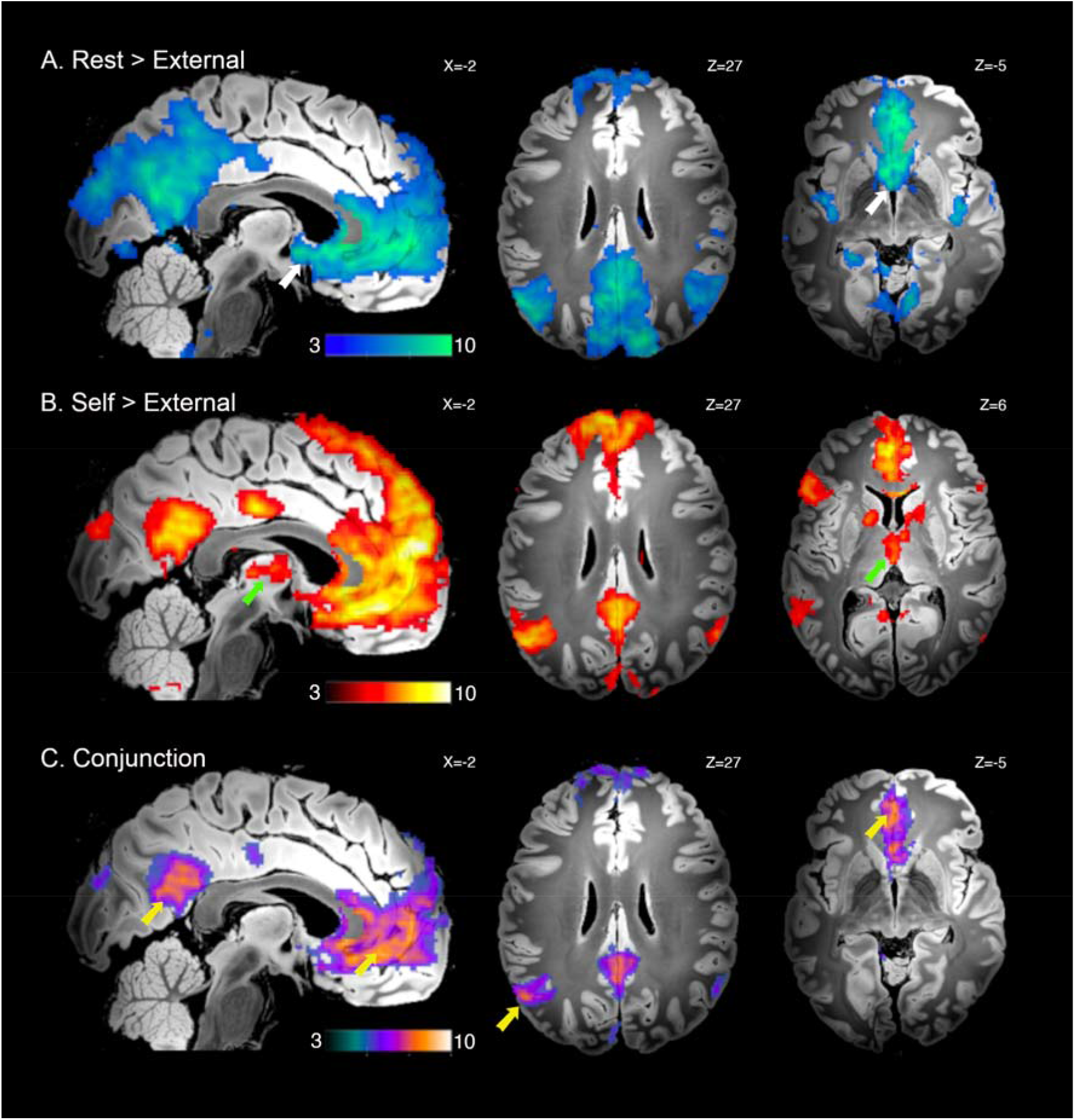
Significant whole-brain UHF fMRI results. A) DMN deactivation, Rest > External conditions; basal forebrain (BF) indicated with white arrows B) DMN activation, Self > External conditions; mediodorsal thalamus (MD) indicated with green arrows C) Conjunction analysis of both A and B; DMN cortical regions indicated with yellow arrows. Displayed contrast maps are thresholded SPM t-statistic images (*P*_FDR_ < 0.05) presented on the ‘Synthesized_FLASH25’ (500um, MNI space) *ex vivo* template (Edlow et al. 2019), with corresponding sagittal and axial slice coordinates.

DMN activation (self > external) was also associated with the broad engagement of characteristic cortical DMN regions: the anterior medial wall cortex, spanning ventral to dorsal MPFC, including rostral anterior cingulate cortex; the posteromedial wall cortex, spanning the PCC, retrosplenial and mid cingulate cortex; the bilateral IPL (angular gyri) and middle temporal gyrus extending to anterior temporal pole (Figure 2B; Table S2). Additional areas of cortical activation included the left frontal opercular, lateral prefrontal and posterior temporal cortices; anterior hippocampus (CA1-3; subiculum); as well as the posterior cerebellum (Crus 1/II). Subcortically, significant activation was observed in the thalamus, encompassing the MD and anterior thalamic nuclei; the medial caudate nucleus (body); the internal globus pallidus; the ventral tegmentum extending to substantia nigra; and the insular claustrum. Activation of the MD was primarily localized to its internal (magnocellular) division, but extended anterior-laterally, reaching the thalamic reticular border (Figure S1B).

The ‘core’ DMN cortical regions identified by conjunction analysis are shown in Figure 2C (Table S3) and confirm a common involvement of the MPFC, with a ventromedial focus; PCC and IPL, particularly the left hemisphere (as in Davey et al. 2016). Additional areas of common engagement included the retrosplenial and mid-cingulate cortex; the medial parieto-occipital zone; the middle temporal gyrus extending to anterior temporal pole; the anterior hippocampus (CA1-3; subiculum); and the posterior cerebellum (Crus 1/II).

Figure S2 provides a further direct comparison between the rest and self conditions. This comparison is useful in illustrating that the BF and MD were significantly differentially engaged across these conditions, with greater BF deactivation occurring during rest versus self, and greater MD activation during self versus rest. All GLM results are presented on the ‘Synthesized_FLASH25’ (500um, MNI space) *ex vivo* template (Edlow et al. 2019).

### Dynamic Causal Modelling

We applied DCM to test specific hypotheses regarding the modulatory influence of the BF (under the rest condition) and MD (under the self condition) on the function of the DMN cortical regions. For the BF and MD, we used inclusive masks from the Alves et al. study to localize group maxima for these regions and to further constrain their time-series extraction across participants. Figure S1 illustrates the specific areas of BF deactivation and MD activation that were localized within the respective boundaries of the Alves et al. masks.

### BF Modulation of the DMN

Under the rest condition, all BF connections, except for the MD-to-BF path, showed evidence of modulation that exceeded our stringent threshold (Pp > 0.99; Figure 3A). BF activity was found to have a strong excitatory (positive) influence on the activity of all cortical DMN regions (range 2.23 to 2.85 Hz), with a low/moderate excitatory influence on MD activity (0.44 Hz). The MPFC had a strong reciprocal excitatory influence on BF activity (2.06 Hz), whereas the PCC (−0.85 Hz) and IPL (−1.16 Hz) had more moderate inhibitory (negative) influences. Table S4 provides a complete list of parameter estimates for the baseline and modulatory connections for the BF model. Compared to the observed modulatory effects, the BF demonstrated low/moderate strength baseline connections with the other regions, and vice versa.

**Figure 3.**
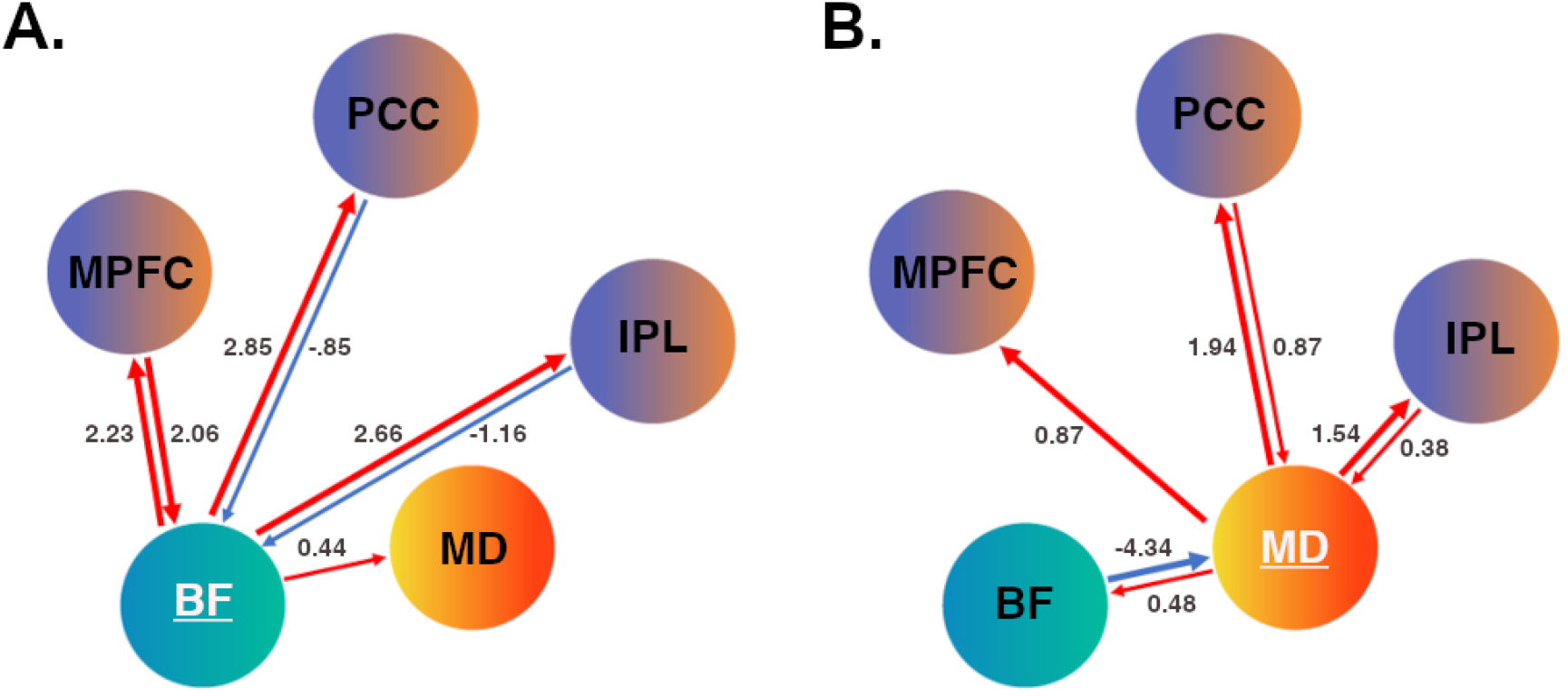
DCM effective connectivity. A) Basal forebrain (BF) modulation under the rest condition; B) Mediodorsal thalamus (MD) modulation under the self condition. Red arrow paths indicate positive/excitatory connections; blue arrow paths indicate negative/inhibitory connections. Respective line thickness indicates strength of connection (≥ or ≤ 1.5Hz). MPFC = medial prefrontal cortex; PCC = posterior cingulate cortex; IPL = inferior parietal lobule.

### MD Modulation of the DMN

Under the self condition, all MD connections, except for the MPFC-to-MD path, showed evidence of modulation that exceeded our threshold (Pp > 0.99; Figure 3B). MD activity was found to have a strong excitatory influence on the activity of all cortical regions (range 0.87 to 1.94 Hz), with a more moderate excitatory influence on BF activity (0.48 Hz). The PCC and IPL had moderate to strong excitatory influence on MD activity (0.87 and −0.38 Hz, respectively), while the BF had a notably strong inhibitory influence (−4.34 Hz). Table S5 provides a complete list of parameter estimates for the baseline and modulatory connections for the MD model. Compared to the observed modulatory effects, the MD demonstrated low/moderate strength baseline connections with other regions (and vice versa), with no significant connections for the MD to BF; PCC or IPL to MD paths. The full set of DCM results (BF and MD models) are available at https://github.com/benharrison-uom/DMN_subcortex_modulation.

### Regional Dynamic Activity

Figure 4 presents group-level (model predicted) responses for the BF (*cyan*), MD (*orange*) and MPFC (*purple*) across the entire task duration and when averaged across a full task epoch (self, rest, external, rest). Figure S3 presents the three cortical regional responses together. These results highlight the consistency of the task evoked neural dynamics, as broadly endorsed by the GLM and DCM findings. Regarding the BF, we note a pattern of evoked responses that preceded changes in MPFC activity during the external and subsequent rest condition blocks, as compared to during the self and subsequent rest condition block. For both the BF and MD, we note a pattern of activity coupling versus decoupling with the MPFC across the self versus external condition blocks, respectively, as well as their more sustained activity across the self condition block.

**Figure 4.**
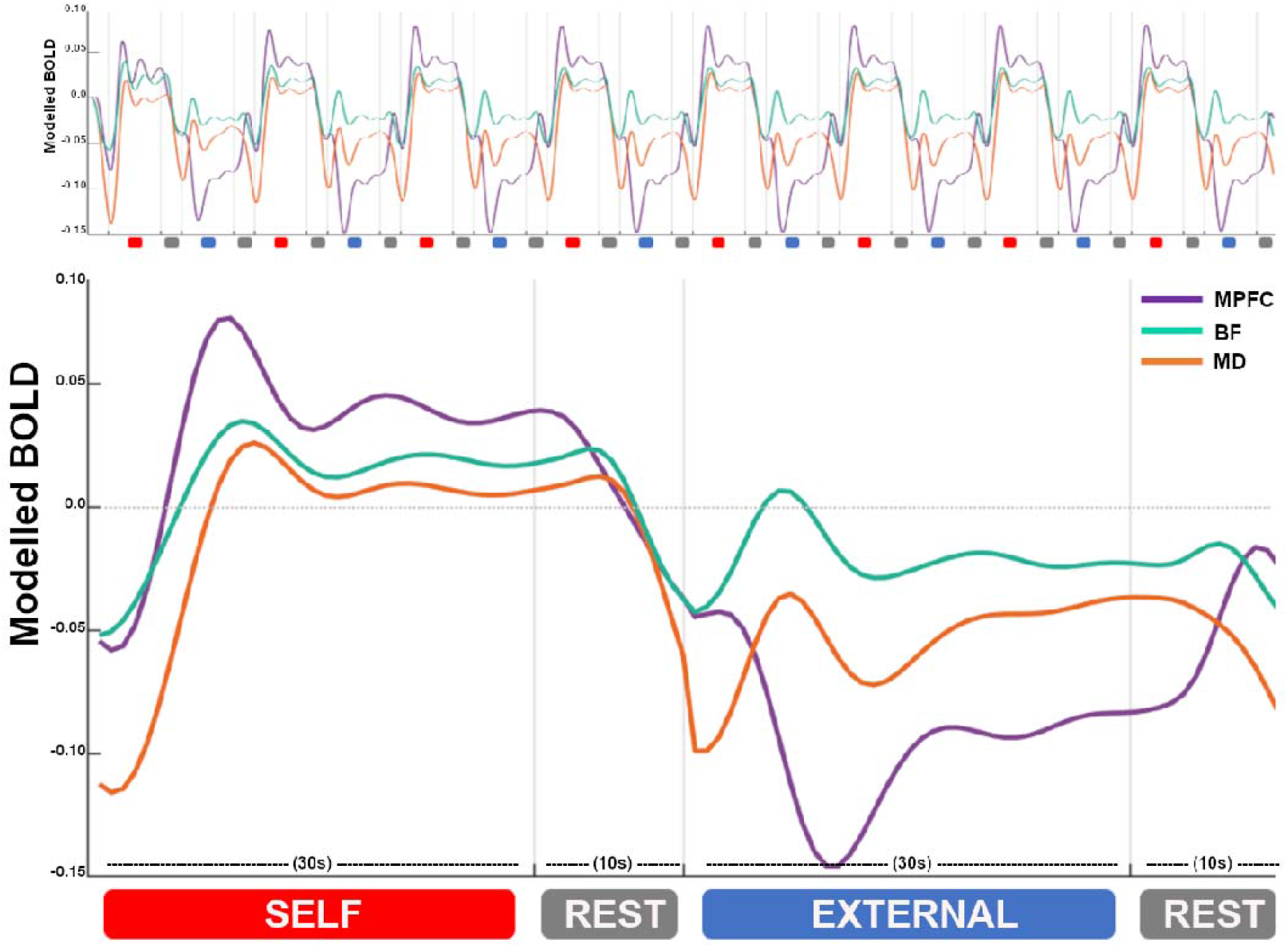
Regional dynamic activity. Group-level (model predicted) responses across the entire task sequence (top) and averaged across all task epochs (bottom). BF responses are represented in cyan; MD responses in orange; and MPFC responses in purple. X-axis = time in seconds (s). Y-axis = estimated BOLD signal change (scaled arbitrary units).

## Discussion

The basal forebrain (BF) and mediodorsal thalamus (MD) have been identified as key subcortical structures of the DMN, with evidence pointing towards their unique contributions to dynamic network functioning (Nair et al. 2018; Alves et al. 2019; Buckner and DiNicola 2019; Steward et al. 2021). In this study, we used task-based UHF fMRI to confirm their involvement across states of DMN activity suppression and engagement, and tested neural systems models of their modulatory influence on DMN cortical regions. Our results endorse a specific role for the BF in driving DMN states transitions between internal and external cognitive modes, extending recent work in rodents (Nair et al. 2018). They also endorse a role for the MD in supporting computations involving major prefrontal circuits (Pergola et al. 2018), which we link here directly to the DMN and its involvement in self-related cognition.

The BF showed an overall pattern of reduced activity during the external task condition compared to rest, suggesting that it was ‘deactivated’ together with the broader DMN, as seen extensively in human functional imaging studies (Shulman et al. 1997; Raichle et al. 2001; Harrison et al. 2008). However, closer inspection of our data revealed that while BF activity was decreased on average when comparing these two conditions, it in fact demonstrated a positive evoked response to the external condition, while DMN cortical regions showed consistent activity suppression (Figures 4/S3). These responses appeared to be driven by the condition’s onset phase (i.e., were not sustained), and consistently preceded changes in DMN cortical regions – an observation that extended to the subsequent rest condition blocks. Given that the self condition, like rest, promotes an internal mental focus, this suggests that BF responses generally tracked the switch between internal and external cognitive states. While DMN cortical regions also exhibited switching responses during the rest to external task transitions, consistent with other recent studies (Crittenden et al. 2015; Smith et al. 2018, 2021), they were less prominent than the BF’s response and were more broadly defined by activity suppression. In the rodent study by Nair et al. (2018), the transition between rest and external task states, especially the reengagement of DMN cortical activity during rest, was causally attributed to BF function. Network modeling of our task’s rest condition also supports these findings: we found the BF to have a robust excitatory influence over DMN cortical regions, whilst receiving distinct reciprocal modulation by the MPFC. We also note that the strength of its influence on DMN cortical regions was several magnitudes stronger that its influence on the MD, which suggests that its influence was likely direct, as opposed to being indirectly mediated via the MD.

In human imaging studies, the BF’s prominent structural and functional connectivity with DMN cortical regions has been suggested to parallel its known modulatory projections involving cholinergic, GABA- and glutamatergic systems (Markello et al. 2018; Alves et al. 2019; Yuan et al. 2019). Using a connectome-based approach, Alves et al. (2019) identified the BF medial septum and diagonal band nuclei as the most densely coupled to DMN cortex, which aligns well with our primary mapping of BF responses. In the Nair et al. study, BF modulation sites more specifically targeted the ventral pallidum and nucleus basalis. Study differences aside, however, the regulation of DMN activity is likely to reflect mechanisms implicating the extended BF circuitry. In recent follow-up studies to Nair et al., for instance, the ascending BF GABAergic (parvalbumin positive; PV+) projection neuron system has been specifically linked to the control of DMN state transitions via its modulation of DMN cortical gamma activity (Lozano-Montes et al. 2020; Klaassen et al. 2021). These neurons, which are distributed throughout the BF in rodents, have long-range connections to DMN cortical regions, particularly with the MPFC and retrosplenial PCC (McKenna et al. 2013; Do et al. 2016). While equivalent data in humans is lacking, there is compelling evidence that links human DMN cortical function to high-gamma modulation (itself a robust correlate of fMRI BOLD signal change; (Fox et al. 2018), as well as local GABA concentrations (Northoff et al. 2007; Hu et al. 2013). Thus, overall, our findings may suggest underlying mechanisms, including GABAergic PV+ influences, that generally index the cortical modulatory function of BF circuitry, particularly in driving cognitive state-transitions.

As expected, the MD was activated with DMN cortical regions during the self compared to the external (and rest) conditions. This result is consistent with past studies, including our own, that have reported significant MD involvement in tasks probing aspects of self-referential cognition (Araujo et al. 2015; Davey et al. 2016). Recently, in Steward et al. (2021), we examined the MD’s specific modulatory influence on distributed prefrontal regions, addressing the hypothesis that it has a key orchestrating influence on the activity of task-relevant higher cortical circuits (Parnaudeau et al. 2018; Pergola et al. 2018; Halassa and Sherman 2019; Wolff and Vann 2019). We used a task that evoked aspects of self-referential cognition and regulatory control and found the MD to have a robust excitatory influence on multiple PFC subregions, whilst receiving distinct reciprocal modulation from the MPFC. Our current results broadly replicate these findings, with the MD again having a robust excitatory influence on DMN cortical regions, while receiving more moderate reciprocal input from the cortex. In Steward et al. (2021), we argued that these broad excitatory influences likely serve to increase the synchrony of cortical regions to help sustain and update complex mental representations (i.e., self representations) that emerge over time. This idea is well supported by animal studies of MD-cortical circuit connectivity (e.g., Schmitt et al. 2017), and gains some further support here, where we observed MD recruitment to be more sustained than DMN cortical regions over the course of the self condition epoch (Figure 4). A focus for ongoing work will be to understand how these dynamics specifically shape patterns of cortico-cortical connectivity across the DMN under distinct cognitive contexts.

We localized MD activity primarily to its internal (magnocellular) segment, which has characteristically dense connections with the MPFC, but only sparse connections with posterior DMN regions (Asanuma et al. 1985; Goldman-Rakic and Porrino 1985; Vogt et al. 1987; Groenewegen 1988; Barbas 2000). Thus, although we observed excitatory links between the MD and DMN cortical regions, the links between the MD and PCC/IPL likely reflect indirect (i.e., intermediary or global network) influences (Adachi et al. 2012). In the MD model, we also identified a prominent inhibitory influence of the BF. The strength of this connection was almost twice that observed for the BF’s excitatory influence on cortical connections (during rest), suggesting that it had a particularly potent influence on MD activity. It is well known that the thalamus receives significant modulatory input from the BF, with the largest proportion of projections (predominately GABAergic) terminating in the reticular nucleus and more moderately in the MD internal segment (Hallanger et al. 1987; Groenewegen 1988; Churchill et al. 1996; Gritti et al. 1998). As noted in Brown and McKenna (2015), these BF projections maintain a high rate of tonic thalamic inhibition, however, in situations requiring cognitive/attentional engagement, their influence is transiently suppressed, potentiates thalamocortical transmission. Importantly, this relationship is recognized to be distinct from the influence of cortically projecting BF GABAergic neurons, which preferentially target inhibitory interneurons, allowing disinhibitory effects and entrainment of fast cortical (i.e., gamma) oscillations. Thus, although fMRI connectivity estimates reflect complex modulatory effects, the observation of distinct cortical excitatory versus thalamic inhibitory effects has a strong anatomical precedent with regards to BF GABAergic function.

## Conclusion

Since its discovery, the study of the DMN has been principally one concerned with higher cortical function, as exemplified by recent perspectives on the DMN as the brain’s ‘apex transmodal association network’ (Margulies et al. 2016; Buckner and DiNicola 2019; Smallwood et al. 2021). However, unique features of the DMN, particularly its characteristic activity suppression during externally focused cognitive tasks, have raised questions about the existence of a subcortical modulatory axis that may support its capacity for dynamic integrative processing. Here, we provide the first direct evidence in humans that confirms distinct basal forebrain and thalamic circuit influences on the control of DMN function. These observations, together with ongoing work, may ultimately help to deliver a more complete, whole-brain perspective on the contribution of the DMN to cognition and behavior, including its role in enigmatic mental processes, such as those related to ‘the self’.

## Notes

We thank Lisa Incerti and Cristian Stella for their contributions to data collection and acknowledge the scientific and technical assistance of the Australian National Imaging Facility – a National Collaborative Research Infrastructure Strategy (NCRIS) capability at the Melbourne Brain Centre Imaging Unit (MBCIU), The University of Melbourne. The multiband fMRI sequence was generously supported by a research collaboration agreement with CMRR, The University of Minnesota and the MP2RAGE works in progress sequence was provided by Siemens Healthineers (Germany).

