## Supplementary data for "Dynamic Subcortical Modulators of Human Default Mode Network Function"

**Contents**

Page 2: Figure S1

Page 3: Figure S2

Page 4: Figure S3

Page 5: Table S1

Page 6: Table S2

Page 7: Table S3

Page 8: Table S4

Page 8: Table S5

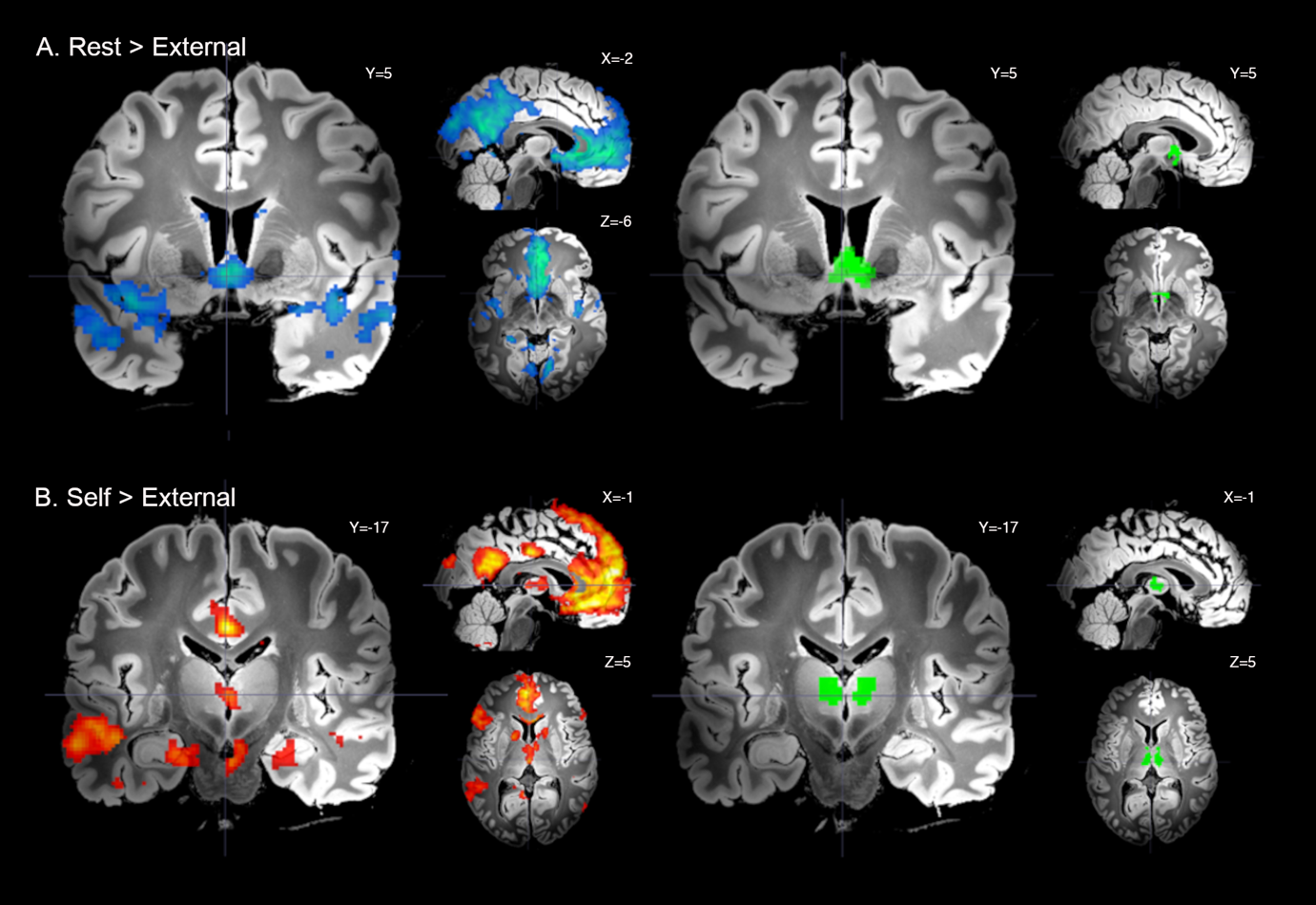

**Figure S1:** Significant whole-brain UHF fMRI results. A) DMN deactivation, Rest > External conditions together with BF mask volume (green) from Alves et al. (2019). B) DMN activation, Self > External condition together with MD mask volume from Alves et al. (2019). Displayed contrast maps are thresholded SPM t-statistic images (*P*_FDR_ < 0.05) presented on the ‘Synthesized_FLASH25’ (500um, MNI space) *ex vivo* template (Edlow et al. 2019), with corresponding sagittal and axial slice coordinates. Crosshairs are included to help localise the primary BF (medial septum/diagonal band) and MD (internal/magnocellular) subregions that were engaged by the task.

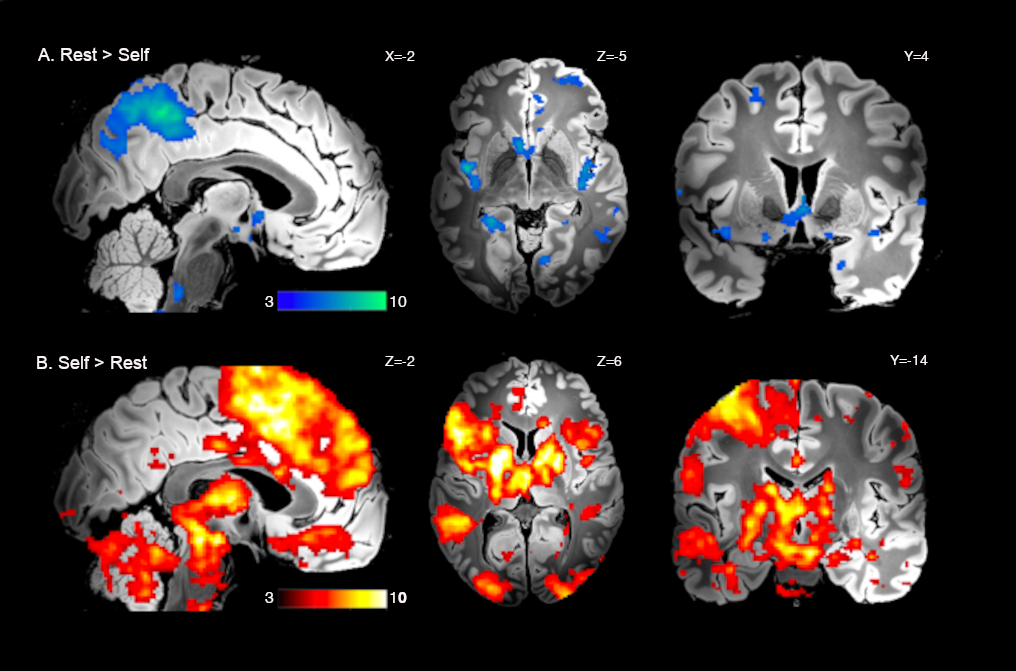

**Figure S2:** Significant whole-brain UHF fMRI results. A) DMN deactivation, Rest > Self conditions; B) DMN activation, Self > Rest conditions. Displayed contrast maps are thresholded SPM t-statistic images (*P*_FDR_ < 0.05) presented on the ‘Synthesized_FLASH25’ (500um, MNI space) *ex vivo* template (Edlow et al. 2019), with corresponding sagittal, axial and coronal slice coordinates.

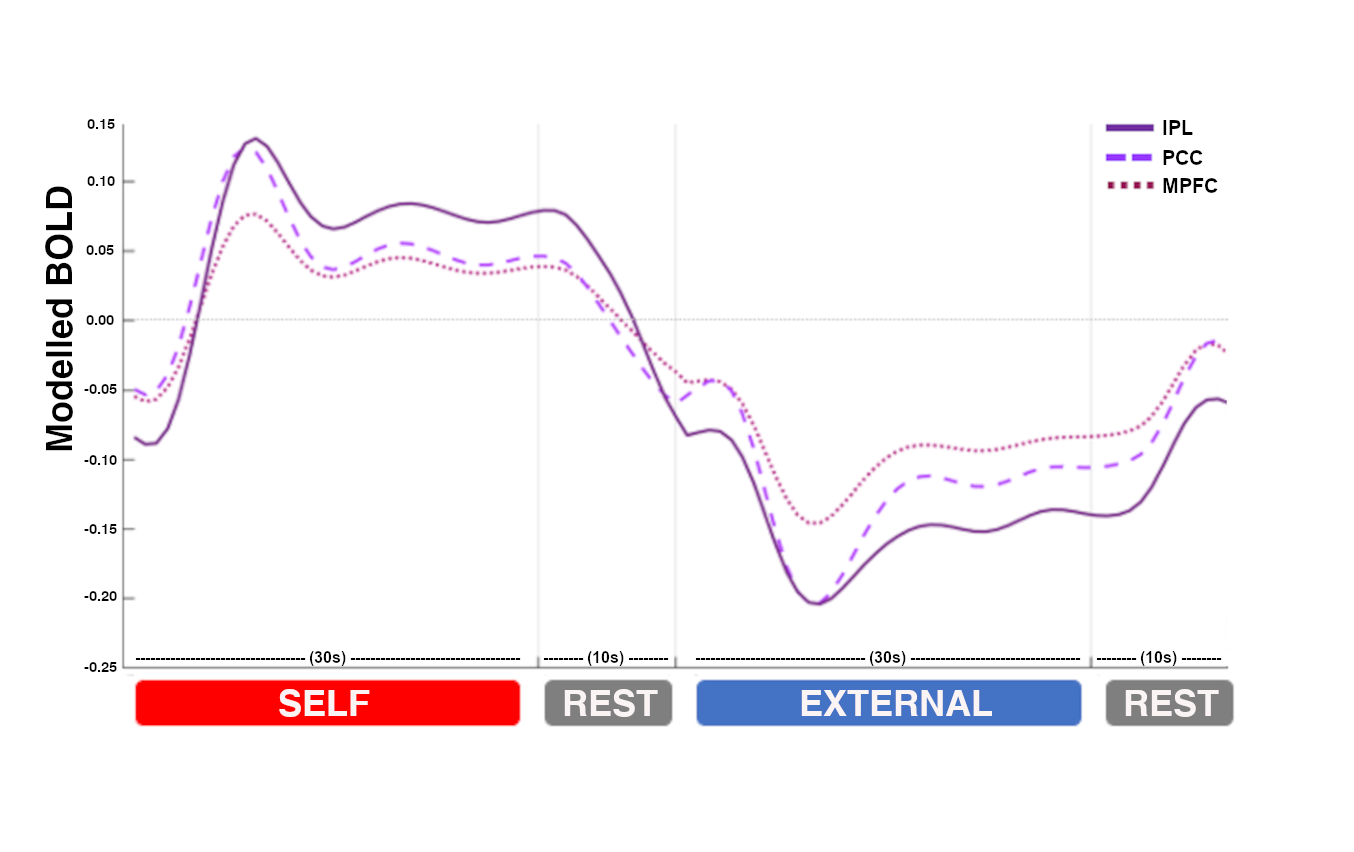

**Figure S3:** Regional dynamic activity. Group-level (model predicted) responses averaged across all task epochs for the three cortical DMN regions. MPFC responses=solid lines; PCC responses=dashed lines; IPL response=dotted lines. X-axis = time in seconds (s). Y-axis = estimated BOLD signal change (scaled arbitrary units).

**Table S1: Significant DMN Deactivation, Rest > External Conditions**

| **Cluster size** | **Brain region** | **SPM Z** | **Peak voxel coordinate** | | |
| --- | --- | --- | --- | --- | --- |
|  |  |  | **x** | **y** | **z** |
| 46168 | Paracingulate gyrus | >8 | 5 | 43 | -6 |
| 3579 | Angular gyrus | 7.68 | -43 | -74 | 35 |
| 8676 | Planum polar | 6.77 | 40 | -19 | -2 |
| 397 | Brain stem | 4.91 | 6 | -45 | -51 |
| 633 | Postcentral gyrus | 4.79 | 50 | -13 | 58 |
| 495 | Cerebellum Crus II | 4.63 | -16 | -80 | -40 |
| 55 | Precentral gyrus | 4.56 | -62 | -2 | 11 |
| 54 | Lateral occipital superior | 4.09 | -24 | -86 | 40 |
| 234 | Anterior insular cortex | 4.07 | -24 | 26 | -8 |
| 17 | Inferior temporal gyrus | 4.07 | 56 | -38 | -22 |
| 63 | Posterior insular cortex | 3.99 | -38 | -40 | 11 |
| 21 | Thalamus (central medial) | 3.94 | 3 | -16 | 2 |
| 40 | Parietal operculum | 3.91 | -53 | -30 | 13 |
| 97 | Inferior frontal gyrus | 3.85 | 59 | 30 | 11 |
| 126 | Cerebellum Crus I | 3.58 | 42 | -75 | -37 |
| 29 | Cerebellum VIII | 3.52 | -13 | -43 | -54 |
| 27 | Precentral SMA | 3.41 | 5 | -14 | 58 |
| 146 | Cerebellum Crus I | 3.41 | 24 | -70 | -35 |
| 98 | Inferior frontal gyrus | 3.4 | 37 | 37 | -6 |
| 25 | Superior frontal gyrus | 3.33 | -10 | 16 | 67 |
| 30 | Temporal pole | 3.33 | 29 | 16 | -35 |
| 10 | Inferior temporal gyrus | 3.3 | -50 | -13 | -32 |
| 27 | Superior frontal gyrus | 3.29 | 0 | 43 | 54 |
| 20 | Medial frontal gyrus | 3.23 | -40 | 16 | 45 |
| 10 | Cerebellum Crus II | 3.2 | 43 | -74 | -42 |
| 22 | Amygdala | 3.2 | 24 | 0 | -32 |
| 19 | Cerebellum Crus II | 3.19 | -45 | -58 | -45 |
| 20 | Brainstem | 3.19 | 5 | -38 | -42 |
| 67 | Cerebellum Crus I | 3.14 | 45 | -53 | -40 |
| 13 | Middle temporal gyrus | 3.11 | -67 | -32 | -2 |
| 12 | Middle temporal gyrus | 3.08 | 66 | -51 | 6 |
| 32 | Pallidum | 3.08 | 22 | 24 | -8 |
| 10 | Paracingulate gyrus | 3.07 | 13 | 38 | 21 |
| 21 | Temporal pole | 3.04 | 38 | 5 | -37 |
| 62 | Cerebellum Crus II | 3.03 | 6 | -86 | -34 |
| 52 | Middle temporal gyrus | 3.03 | 64 | -42 | 2 |
| 23 | Middle temporal gyrus | 3.02 | -64 | -54 | 5 |
| 13 | Thalamus (central medial) | 2.91 | 8 | -30 | 2 |
| 19 | Postcentral gyrus | 2.91 | -38 | -19 | 35 |
| 13 | Lateral occipital cortex | 2.9 | -54 | -70 | 5 |

Cluster sizes = # of voxels; SPM Z = Z scores; Coordinates are reported in MNI space.

**Table S2: Significant DMN Activation, Self > External Conditions**

| **Cluster size** | **Brain region** | **SPM Z** | **Peak voxel coordinate** | | |
| --- | --- | --- | --- | --- | --- |
|  |  |  | **x** | **y** | **z** |
| 22101 | Superior frontal gyrus | 7.76 | -13 | - 53 | 42 |
| 3007 | Cerebellum Crus I/II | 7.71 | 27 | -78 | -35 |
| 19843 | Middle temporal gyrus | 7.12 | -46 | 5 | -30 |
| 418 | Cingulate gyrus | 6.48 | 0 | -18 | 34 |
| 763 | Cerebellum Crus I/II | 5.97 | -26 | -80 | -32 |
| 2788 | Middle temporal gyrus | 5.52 | 48 | 13 | -30 |
| 387 | Brainstem | 5.45 | 6 | -43 | -48 |
| 610 | Inferior frontal gyrus | 4.73 | 38 | 29 | -13 |
| 535 | Angular gyrus | 4.68 | 58 | -62 | 19 |
| 30 | Pallidum | 4.36 | -22 | -13 | -6 |
| 295 | Occipital pole | 4.02 | -2 | -88 | 24 |
| 56 | Posterior insular cortex | 3.92 | -40 | -2 | -16 |
| 28 | Parahippocampal gyrus | 3.84 | 18 | -32 | -19 |
| 52 | Midbrain (~substantia nigra) | 3.76 | -8 | -22 | -13 |
| 30 | Lateral occipital cortex | 3.66 | 58 | -64 | 3 |
| 184 | Occipital pole | 3.42 | 21 | -96 | 26 |
| 13 | Hippocampus | 3.29 | 34 | -38 | -6 |
| 19 | Parahippocampal gyrus | 3.23 | 24 | -2 | -34 |
| 74 | Middle temporal gyrus | 3.17 | 46 | -37 | 0 |
| 34 | Inferior frontal gyrus | 3.17 | 58 | 35 | 13 |
| 11 | Lingual gyrus | 2.96 | 11 | -50 | 2 |
| 15 | Cingulate gyrus | 2.94 | -14 | -45 | 6 |
| 11 | Lingual gyrus | 2.82 | 13 | -69 | -8 |

Cluster sizes = # of voxels; SPM Z = Z scores; Coordinates are reported in MNI space.

**Table S3: Significant Conjunction (Rest > External AND Self > External conditions)**

| **Cluster size** | **Brain region** | **SPM Z** | **Peak voxel coordinate** | | |
| --- | --- | --- | --- | --- | --- |
|  |  |  | **x** | **y** | **z** |
| 8963 | Paracingulate gyrus | 6.89 | 2 | 40 | -8 |
| 2261 | Cingulate gyrus | 6.3 | 2 | -43 | 24 |
| 1602 | Angular gyrus | 5.76 | -53 | -66 | 27 |
| 1441 | Middle temporal gyrus | 5.42 | -64 | -5 | -21 |
| 753 | Middle temporal gyrus | 4.91 | 56 | -5 | -16 |
| 194 | Cingulate gyrus | 4.64 | 2 | -18 | 34 |
| 95 | Brainstem | 4.51 | 6 | -45 | -51 |
| 111 | Insular cortex | 4.49 | -32 | 8 | -11 |
| 313 | Angular gyrus | 4.45 | 58 | -62 | 19 |
| 124 | Hippocampus | 4.39 | 27 | -10 | -21 |
| 227 | Cerebellum Crus II | 4.33 | -18 | -80 | -40 |
| 142 | Insular cortex | 4.26 | 32 | 13 | -16 |
| 214 | Hippocampus | 4.17 | -21 | -22 | -19 |
| 107 | Cuneus | 4.02 | -2 | -88 | 24 |
| 11 | Insular cortex | 3.92 | -40 | -2 | -16 |
| 17 | Parahippocampal gyrus | 3.84 | 18 | -32 | -19 |
| 50 | Superior frontal gyrus | 3.73 | 19 | 32 | 54 |
| 50 | Middle frontal gyrus | 3.68 | -35 | 24 | 51 |
| 41 | Inferior frontal gyrus | 3.56 | -30 | 29 | -13 |
| 22 | Parahippocampal gyrus | 3.54 | 14 | -11 | -26 |
| 35 | Cerebellum Crus I | 3.41 | 24 | -70 | -35 |
| 18 | Frontal pole | 3.34 | -22 | 48 | 26 |
| 11 | Superior frontal gyrus | 3.33 | -10 | 16 | 67 |
| 31 | Occipital pole | 3.28 | 14 | -98 | 22 |

Cluster sizes = # of voxels; SPM Z = Z scores; Coordinates are reported in MNI space.

**Table S4: ‘BF-Rest’ Model Parameter Estimates**

| **Connection** | **Strength (Hz)** | **Covariance** |
| --- | --- | --- |
| **Baseline effects** | |  |
| BF → MPFC | -0.34 | 0.0006 |
| BF → PCC | -0.57 | 0.0009 |
| BF → IPL | -0.19 | 0.0008 |
| BF → MD | 0.19 | 0.0005 |
| MPFC → BF | 0.26 | 0.0009 |
| PCC → BF | -0.16 | 0.0005 |
| IPL → BF | 0.18 | 0.0004 |
| MD → BF | -0.28 | 0.0005 |
| **Modulatory effects** | |  |
| BF → MPFC | 2.23 | 0.0122 |
| BF → PCC | 2.86 | 0.0135 |
| BF → IPL | 2.67 | 0.0215 |
| BF → MD | 0.45 | 0.0068 |
| MPFC → BF | 2.07 | 0.0166 |
| PCC → BF | -0.85 | 0.0094 |
| IPL → BF | -1.16 | 0.0087 |
| MD → BF | - | - |

*Estimated strength in Hertz (Hz) and uncertainty (i.e., covariance) of significant connections (posterior probability > .99). BF=Basal Forebrain; MPFC=Medial Frontal Cortex; PCC=Posterior Cingulate Cortex; IPL=Inferior Parietal Lobule; MD=Mediodorsal Thalamus.*

**Table S5: ‘MD-Self’ Model Parameter Estimates**

| **Connection** | **Strength (Hz)** |  |
| --- | --- | --- |
| **Baseline effects** | |  |
| MD → MPFC | -0.09 | 0.0004 |
| MD → PCC | -0.12 | 0.0007 |
| MD → IPL | 0.23 | 0.0006 |
| MD → BF | - | - |
| MPFC → MD | 0.13 | 0.0012 |
| PCC → MD | - | - |
| IPL → MD | - | - |
| BF → MD | 0.29 | 0.002 |
| **Modulatory effects** | |  |
| MD → MPFC | 0.87 | 0.0074 |
| MD → PCC | 1.94 | 0.0099 |
| MD → IPL | 1.54 | 0.0077 |
| MD → BF | 0.48 | 0.0027 |
| MPFC → MD | - | - |
| PCC → MD | 0.87 | 0.0078 |
| IPL → MD | 0.38 | 0.0057 |
| BF → MD | -4.34 | 0.0388 |

*Estimated strength in Hertz (Hz) and uncertainty (i.e., covariance) of significant connections (posterior probability > .99). BF=Basal Forebrain; MPFC=Medial Frontal Cortex; PCC=Posterior Cingulate Cortex; IPL=Inferior Parietal Lobule; MD=Mediodorsal Thalamus.*
